# A trigger may not be necessary to cause senescence in deciduous broadleaf forests

**DOI:** 10.1101/2023.06.07.544057

**Authors:** Kathryn I. Wheeler, Michael C. Dietze

## Abstract

Plant phenological changes drive many ecosystem processes and are a key ecological indicator of climate change. Traditional models represent the onset of autumn leaf senescence, or color change, as a threshold response triggered by the accumulation of cold temperatures and declining day lengths, but the physiological mechanisms behind plant thermal memory and chilling thresholds remain elusive. Here we show that we can predict senescence in forest canopies by dynamically modeling daily greenness as the “memoryless” balance between chlorophyll synthesis, linearly-related to daily temperatures and day lengths, and breakdown. Indeed, summer-only data can be used to estimate breakdown and synthesis rates that in many cases successfully predict senescence at both calibration and validation sites. This mechanistic model shows that neither a trigger nor a physiological memory of coldness is necessary for senescence to start. These findings suggest that the start of senescence is not an irreversible transition, but a continuum of decreasing greenness where concurrent environmental conditions determine the rate of initial senescence. Furthermore, this emphasizes that in order to predict how senescence will shift in response to global change we likely need to focus on understanding the impacts on chlorophyll synthesis.

**Significance statement:** Plant phenology is a leading ecological indicator of climate change and has wide-ranging ecological and climatological impacts. Our findings here contradict the key assumption that senescence in deciduous broadleaf trees is actively triggered only when a threshold of cumulative cooling in combination of other stressors is reached. Instead we show that canopy greenness can be modeled as a passive process – balancing chlorophyll breakdown and temperature- and photoperiod-controlled synthesis – and still predict a rapid decline in greenness during senescence. This work is particularly important because it emphasizes that in order to understand climate change induced shifts in senescence, we need to focus on impacts on chlorophyll synthesis.

## 1. Introduction

Phenology is one of the primary ecological indicators of climate change (1–4). Due to the numerous controls that vegetation phenology exerts on ecosystems, climate change’s lengthening of the growing season impacts diverse ecosystem processes such as annual primary productivity, energy budgets, biogeochemical cycling, and weather (5–9). Within cold-deciduous broadleaf (DB) forests, autumn senescence (rapid decrease in greenness) is generally understudied, less understood, and harder to predict than spring (2). Senescence is important for plants, as chlorophyll and nutrient concentrations decrease in preparation for dormancy, and temperature and photoperiod (*i.e.,* day length) are the primary drivers of senescence (2). The impacts of climate change, which alters temperatures but not photoperiod, varies between sites: photoperiod is generally dominant in areas with harsh winters, while temperature is more important in sites with milder winters (10).

Despite this general understanding, current models are inadequate at predicting the end of the growing season (6) and senescence under novel conditions, either at other sites or under climate change (11). Most phenology models represent statistical relationships between environmental drivers and the “event” timing of transition dates (12–17), rather than the dynamic ecophysiological process(es) underlying senescence (11), and, thus, face limitations when predicting under future, novel conditions. At the canopy level, senescence is traditionally modeled as a trigger, where once a certain cooling degree day threshold (CDD; *i.e.,* the cumulative amount that daily temperature is below a reference value) or photoperiod is reached, senescence proceeds irreversibly. CDD models, however, are problematic. They are sensitive to the subjective selection of base temperatures and start dates and provide limited understanding of what is happening physiologically (18). They assume there are mechanisms by which plants sense thermal sums, accumulate them over time, and store that information. Some suggest that a potential mechanism for this could be accumulation of sugars and starches, but this is still unclear (19, 20). The need for a senescence trigger has been questioned before (21), but it has not been investigated and is assumed in models. Developing more mechanistic models of senescence requires that we critically reexamine these assumptions (11).

At a fundamental level, senescence is a visible rapid decrease in leaf chlorophyll concentrations. However even in the growing season, chlorophyll is constantly cycled via breakdown and synthesis (22). Consistent with observations of the drivers of senescence, chlorophyll synthesis is temperature dependent, as a product of enzyme-mediated reactions, and is likely limited to the daytime in angiosperms (23, 24). Therefore, chlorophyll cycling can serve as a framework for modeling senescence as a “memoryless” dynamic process (*i.e.,* condition at *time = t* + 1 depends on the condition at *time = t*), based on the balance between chlorophyll synthesis and breakdown.

We postulate that senescence in DB trees results from a temperature- and photoperiod-driven decrease in synthesis, relative to breakdown, and not because of an inherent trigger when a threshold of accumulated coldness or day length is reached. While a trigger-based approach might be supported by the internal transitions from anabolic to catabolic genes necessary to facilitate the resorption of key nutrients from the leaves and prevent late autumn increases in chlorophyll concentrations (25), this does not necessarily mean that the start of senescence (*i.e.,* the inflection where greenness starts to decrease more sharply; SOS) is an irreversible physiological transition. Senescence is different from programmed cell death because of its reversibility (21, 26–28), which manifests in canopy regreening events in the late growing season and at the cellular level where gerontoplasts, which are an intermediate product of chlorophyll breakdown, are able to re-differentiate back into green chloroplasts (29, 30). Indeed, the transition to a catabolic state may be responding to, rather than causing, senescence.

In support of the hypothesis that SOS is a continuous process, rather than a trigger, is the observation that the rate at which canopy greenness is lost at the SOS can vary considerably between sites and from year-to-year. Since conventional CDD approaches are trying to predict threshold dates, these models do not distinguish between sharp and gradual SOS and require historical SOS for calibration. However, if SOS is governed by the same chlorophyll synthesis and breakdown processes as occur in the growing season, then one could theoretically (a) predict post-SOS greenness through calibrating the model to solely data within the growing season (pre-SOS) and (b) predict the amount of curvature in SOS. Doing so would represent a strong test of this mechanistic framework that has not been done elsewhere.

Remote sensing data, such as PhenoCam digital cameras, provide information about the full autumn GD curve, rather than just discrete transition dates, and, thus, enable the development of dynamic mechanistic models. PhenoCam greenness observations for DB systems typically observe a gradual decline in chlorophyll across the summer, followed by leaf senescence, and finally abscission (*i.e.,* leaf drop; Fig. 1). Here we use GD to refer to all three processes and senescence to just refer to the rapid leaf color change right after an inflection.

**Fig. 1.**
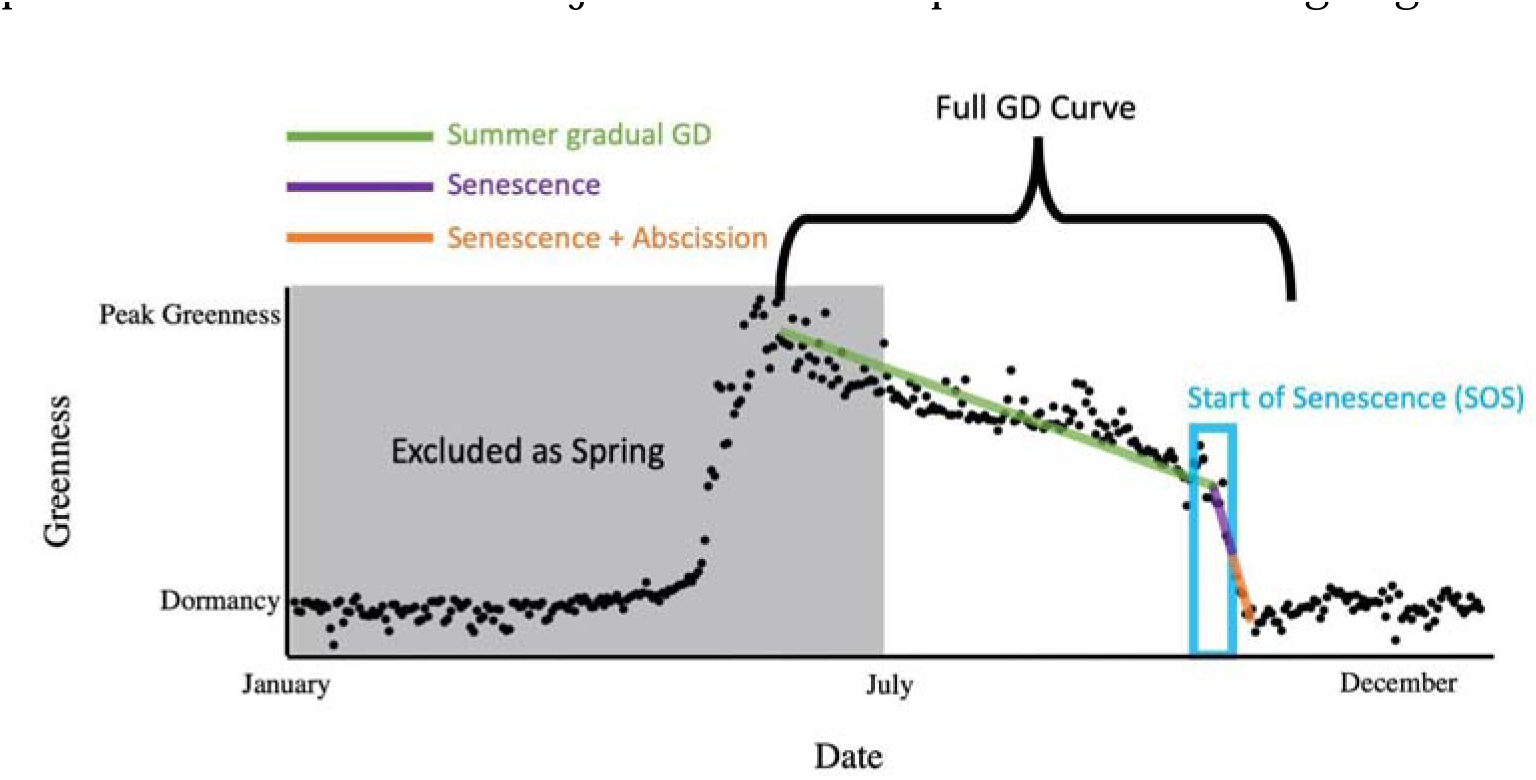
Parts of the yearly greenness curve indicated. Gray shading indicates data we excluded as spring, which was defined as 1 January through 30 June of each year. Full green-down (GD) encompasses the summer gradual GD, senescence, and abscission. In this manuscript we were particularly interested in predicting the greenness and amount of inflection around the SOS (blue box).

In this study we predicted daily autumn greenness at seventy DB PhenoCam sites (Supplementary Table S1) by modeling the continuous cycling of greenness breakdown and synthesis, where synthesis is linearly related to the product of temperature and photoperiod and breakdown follows a constant exponential decay. We first calibrated the model individually at 24 PhenoCam sites to the full autumn. To investigate whether growing season chlorophyll dynamics could predict senescence, we then iteratively recalibrated the models with one less day of year from the end of the timeseries until model convergence was no longer achievable. We then used the calibrated models for each of these sites to predict greenness at other sites. The model fits were compared to predictions based on each site’s historical average GD, which is typically hard for phenology models to outperform (31) because the SOS timing is often similar year-to-year with a strong photoperiod control (11).

Our overarching hypothesis was that the SOS in DB is often due to a passive process of declining chlorophyll synthesis and not an active trigger. If this is true, then we further hypothesized (H1) a chlorophyll-cycling model without a CDD threshold-based trigger performs better than historical predictions around, and right after, the SOS for calibration and validation sites; and (H2) We can estimate the synthesis and breakdown parameters necessary to cause rapid declines of greenness during senescence and the variation in the amount of curvature of the SOS using only pre-senescence calibration data.

To provide further support for the overarching hypothesis independent of our model selection, we also investigated if the air temperatures and photoperiods before or after a date had more power in predicting SOS occurrence using a random forest model. If greenness is more rapidly declining after SOS because the concurrent air temperatures and photoperiods limit chlorophyll synthesis and not because of conditions before triggering SOS, (H3) environmental drivers after a date would be better predictors of if SOS occurred than the environmental drivers before the date.

## 2. Results

### 2.1 Example chlorophyll cycling model fits

In support of our overarching hypothesis, the chlorophyll cycling model even though it did not have a trigger to create a SOS inflection and a rapid decline in greenness was able to fit to all of our calibration sites and predict inflections in the withheld year (example fits given in Fig. 2). For sites like Howland, Maine, USA, that have a moderate amount of noise in PhenoCam observations, the within-site validation year’s credible interval (CI), which is the Bayesian equivalent of the confidence interval, was wide, but still predicted a dramatic decrease in greenness within days of the PhenoCam SOS inflection (Fig. 2a). Sites with less day-to-day variation in PhenoCam observations, such as Alligator River, North Carolina, USA, had much narrower predictions for the validation year (Fig. 2b). Furthermore, there is evidence that our model predicts regreening events (Fig. 2e). Calibrated parameters for all days at each site are given in Supplementary Table S2.

**Fig. 2.**
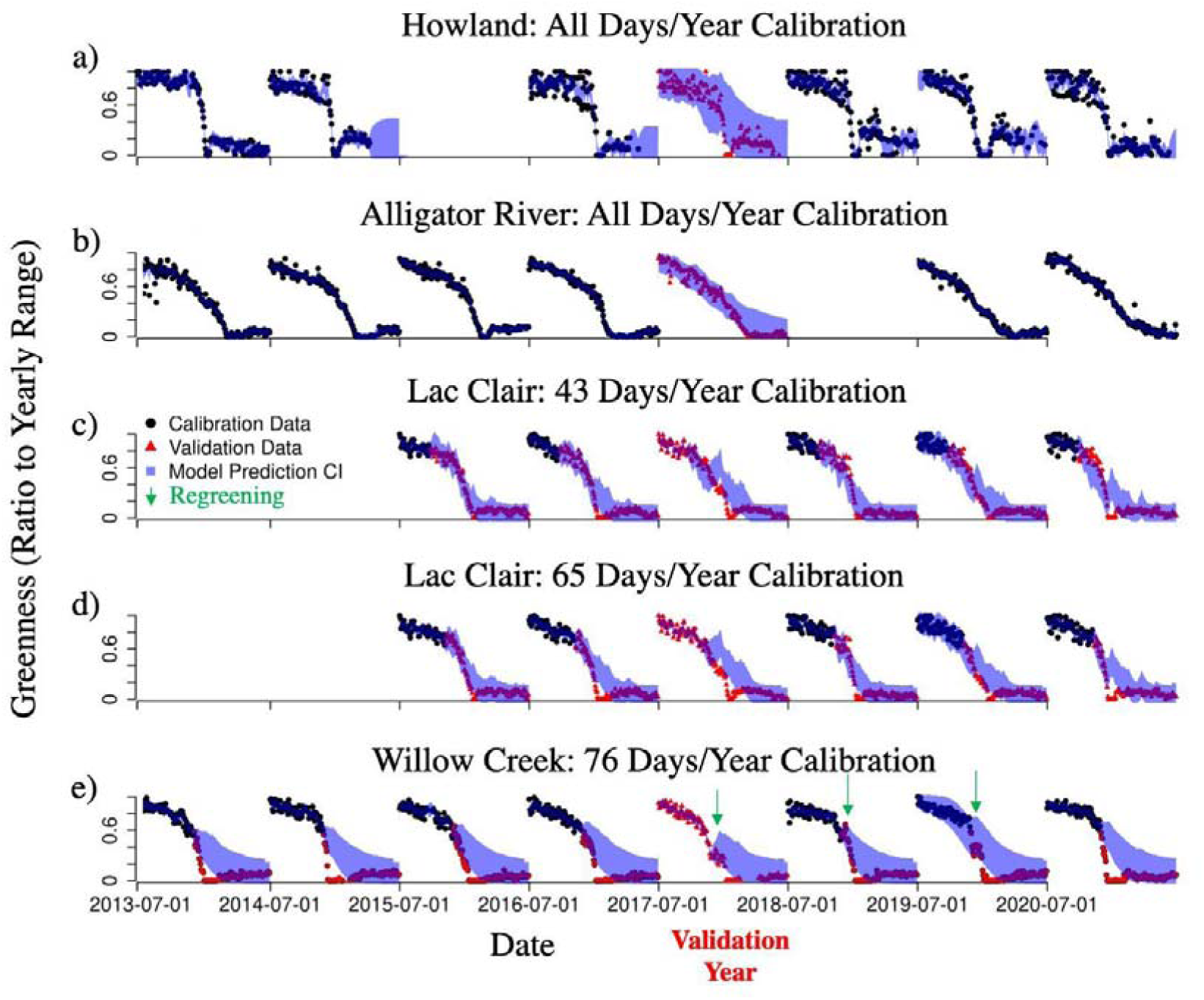
Example time series of modeled 95% credible intervals (blue shading) and calibration (black circles) and validation (red triangle) data for (a) Howland, Maine, USA, calibrated to the full autumn(1 July – 31 December); (b) Alligator River, North Carolina, USA, calibrated to full autumn; Lac Clair, Quebec, Canada calibrated to 43 days per year (c) versus 65 days per year (d); and (e) Willow Creek, Wisconsin, USA calibrated to 76 days per year. The model was able to produce reasonable estimates of greenness for withheld data even without including any senescence greenness data and potentially predicted regreening events (green arrows point to potential observed re-greening).

### 2.2. H1: Predicting Greenness Around Start of Senescence Better Than Historical Average

In support of H1, our model predicted greenness around the SOS and throughout the autumn better than historical averages in all the site-years and the withheld validation years of the calibration sites (Fig. 3). Since smaller values are better, negative CRPS differences (historical average’s CRPS - our model’s CRPS) indicated that our model performed better than the historical averages estimate for both calibration and validation years (Fig. 3). The strongest improvement in predictability compared to historical prediction occurred around 20 days post-SOS.

**Fig. 3.**
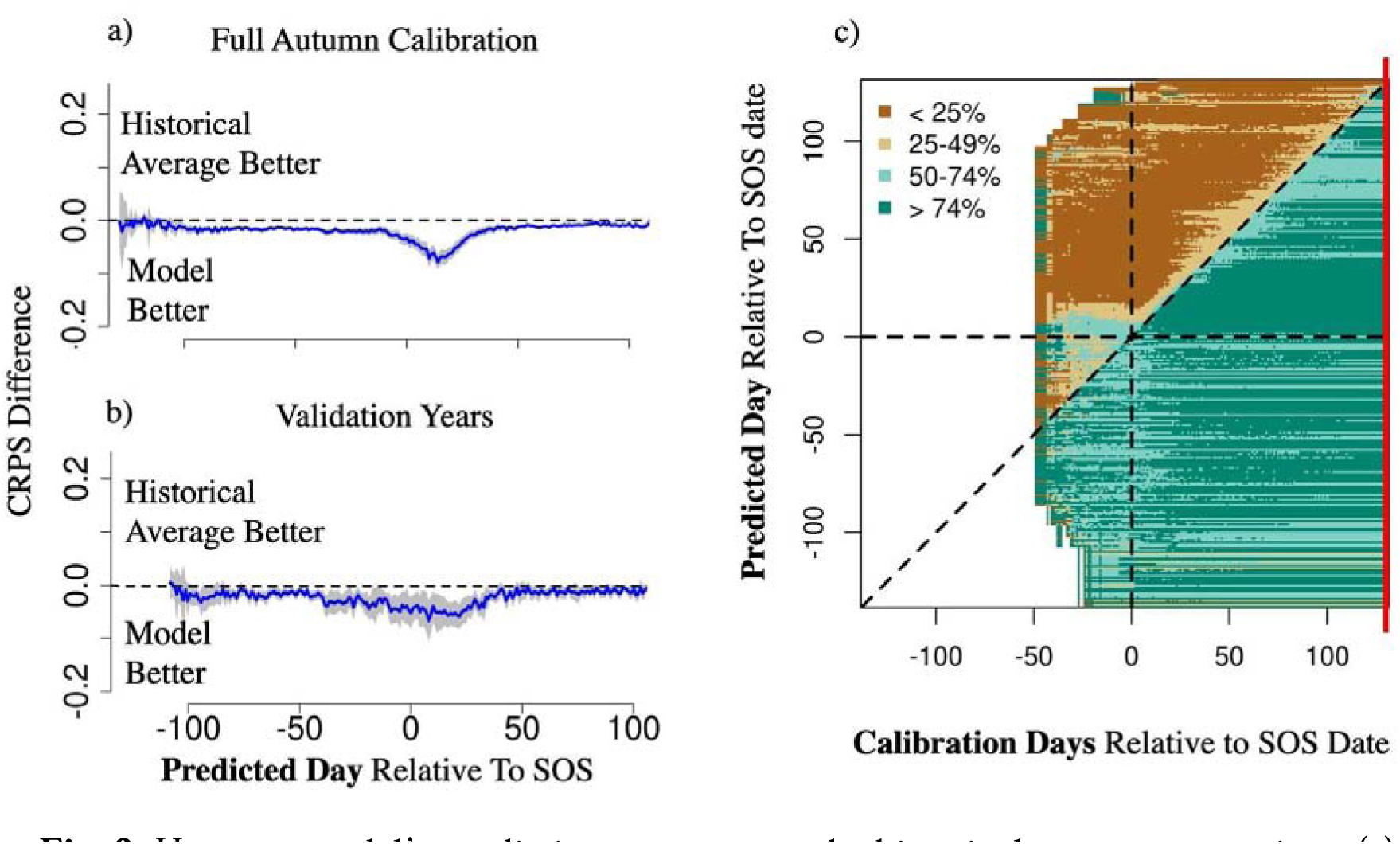
How our model’s predictions compare to the historical averages over time. (a) Continuous Ranked Probability Score (CRPS) differences over time (mean dark blue line and standard error in gray shading) across all site-years relative to start of senescence (SOS) date when the models were calibrated to the entire autumn. (b) CRPS differences over time across validation years. (c) Heat map showing the percentage of site-years that our model was better at predicting specific days (relative to SOS) as a function of the days the model was calibrated to. The vertical, solid, red line in (c) shows where the example time-series from (a) occurs. The model consistently and significantly predicted greenness at the beginning of senescence better than climatology even when it was not calibrated to any senescence data.

In further support of H1, models calibrated to some sites (Supplementary Table S3) had high transferability to other sites (*i.e.,* had lower CRPS values than the validation site’s historical averages for different periods in the autumn; Fig. S1). The transferability was higher for the periods that model performed best in the calibration sites (*e.g.,* six days after the SOS > within three days > six days before) and lowest for the full autumn (Fig. S1). The factors that had the largest influence on the transferability were the differences in mean annual temperature (MAT; *p*-value: 0.000578), maximum yearly PhenoCam greenness (*p-*value: 0.016255), and latitude (*p-* value: 0.040688) between calibration and validation sites. In all cases transferability was higher when the differences between sites was smaller. Other tested explanatory variables (Supplementary Table S1) were not significantly correlated with transferability.

### 2.3. H2: Synthesis and breakdown parameters can be estimated from only pre-senescence data

In support of H2, model calibrations that only included data prior to SOS successfully predicted an inflection in greenness values similar to what constitutes a SOS inflection in PhenoCam data in 49% and 46% of the calibration and validation site-years, respectively, without including any senescence data in the calibration (Fig. S2). The model significantly explained variation in the amount of inflection between site-years when calibrated to only include through one day before SOS data (Fig. 4).

**Fig. 4.**
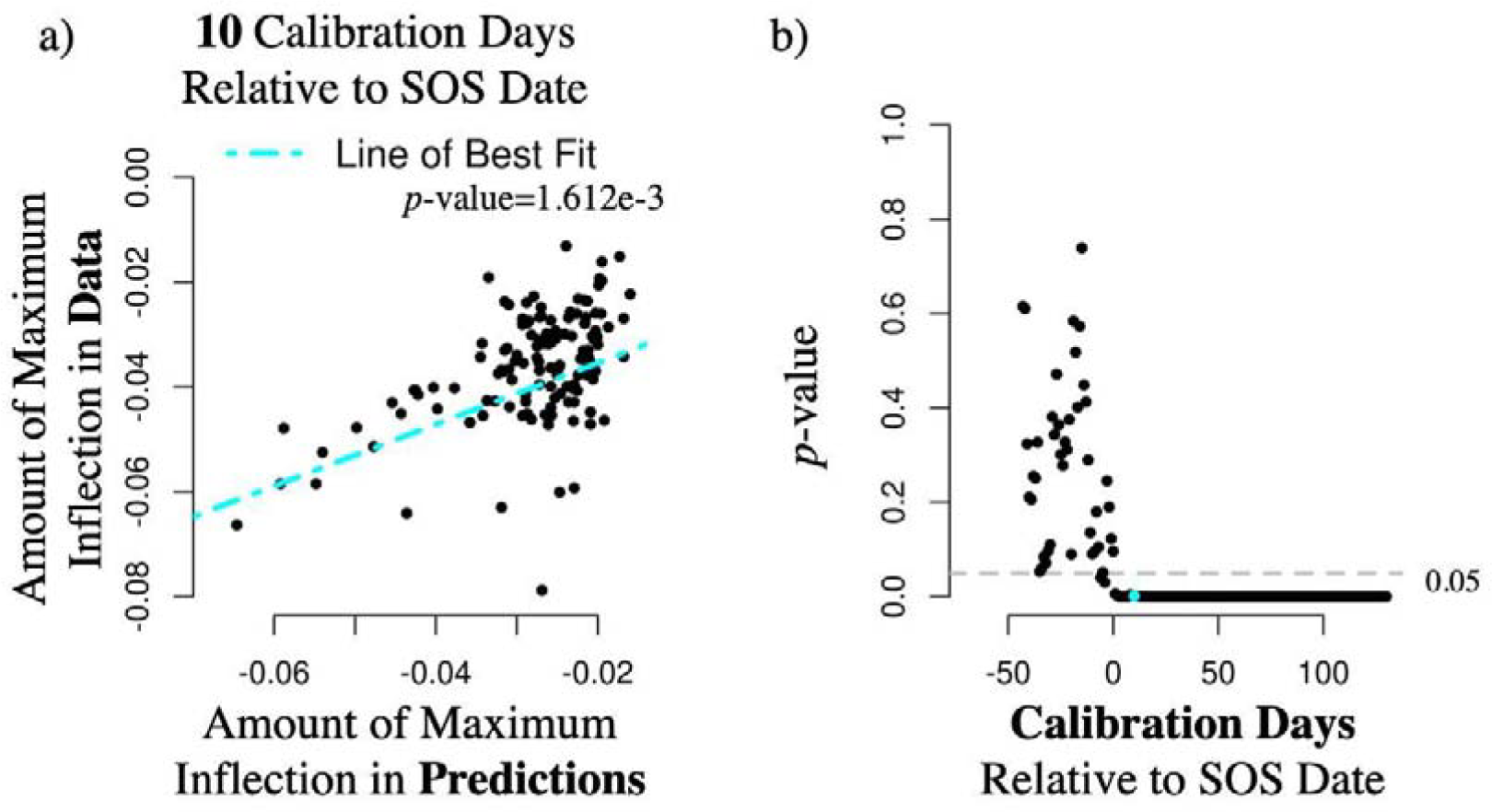
The ability of our model to predict significant variation in the amount of curvature of start of senescence (SOS). (a) Example regression between the predicted amount of inflection when only calibrating through 10 days after SOS compared to the amount of maximum inflection seen in the PhenoCam data. (b) Linear regression *p-*values comparing data maximum inflection vs predicted maximum inflection as a function of how much data is included in the calibration relative to the SOS date. The blue point in (b) indicates the p-value from example (a). Accompanying statistics are given in Fig. S3. Our model consistently predicted variation across calibration site-years in the sharpness of the SOS inflection when only including through 1 day before the average SOS date.

For the majority of these site-years, the “summer-only” calibrations of the model also predicted greenness around the SOS (+/-10 days) better than historical averages (Fig. 3a, c). For example, calibrations through 0–25 days before SOS produced greenness estimates that were better than historical averages for the week after SOS in an average of 61% of site-years (standard deviation=0.056). When including various amounts of days relative to the SOS the majority of all site-years performed better than historical averages for those predicted days and earlier (Fig. 3c).

For example, at Lac Clair, Quebec, Canada (Fig. 2c) the model predicted reasonable greenness estimates for the entire season despite only being calibrated against PhenoCam data from 1 July to 12 August (43 days per year), including capturing the delayed SOS in 2015 and the early and less pronounced inflections in 2019 and 2020. By including more days (for example, through approximately 4 September) each year, the model more accurately and precisely captured the sharp GD in 2020 (Fig. 2d), which was predicted slightly earlier when the model was calibrated with only 43 days per year.

### 2.4. H3: Drivers After Date Better Predictor of SOS Than Those Before

In further support of our hypothesis that a trigger is not necessary for SOS and in support of H3, our random forest analysis indicated that most important factor for predicting if the SOS inflection occurred or not was the temperature × day length interaction the week after the date being predicted (Fig. 5). The importance of the other predictors, in decreasing order, were temperature × day length before, day length after, day length before, temperature after, temperature before, latitude, and mean annual temperature. Across each of the environmental drivers, the mean value the week after a date was a more important predictor in the model of SOS than the mean value the week before a date.

**Fig. 5.**
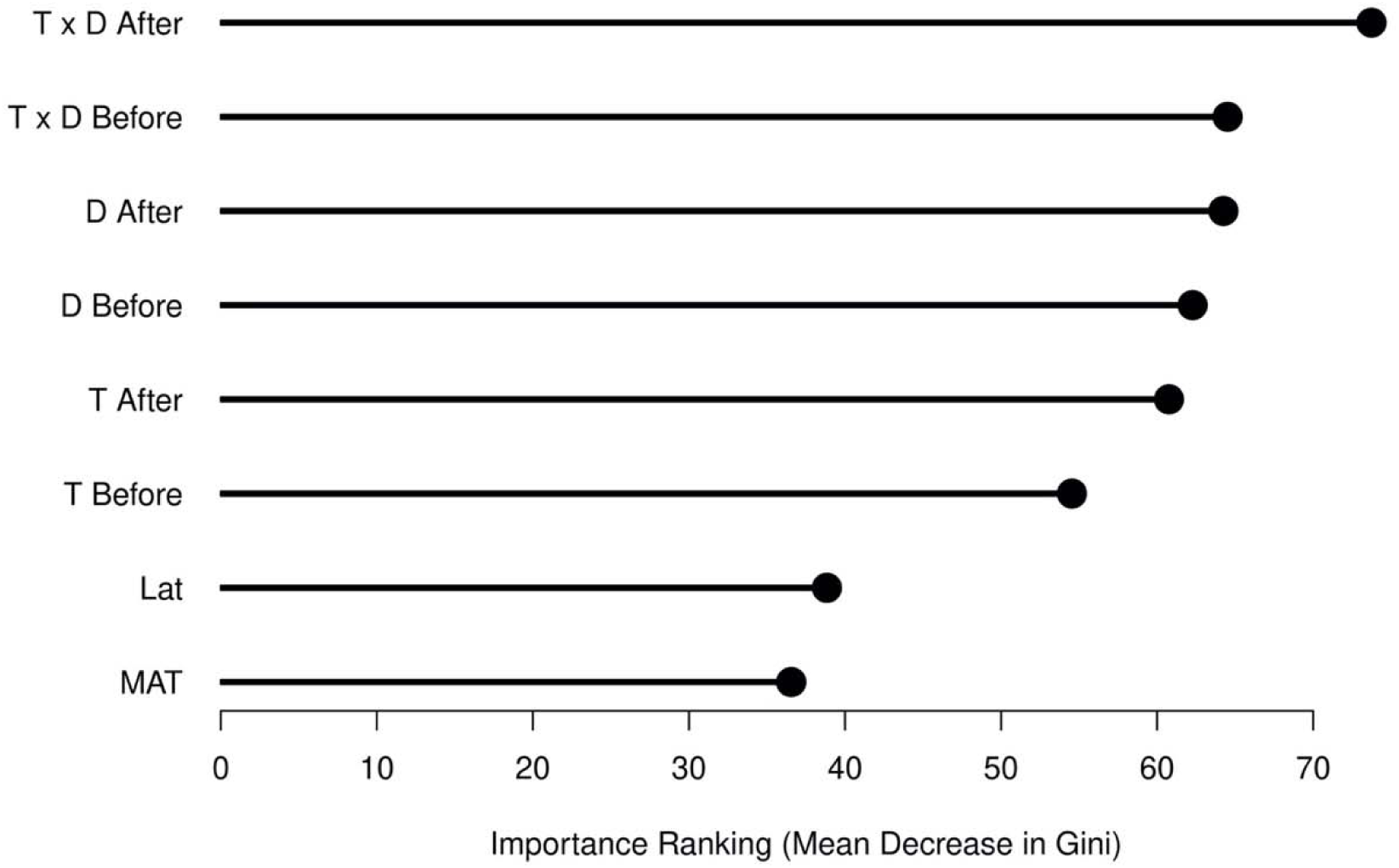
Importance ranking of predictors in the fitted random forest model to predict if the start of senescence inflection occurred or not. The mean product of temperature and day length the week after a date was the most important predictor.

## 3. Discussion

### 3.1. H1: Predicting Greenness Around Start of Senescence Better Than Historical Average

Our results support our hypothesis that we can predict greenness around the SOS better than historical averages without including a trigger, which has numerous implications for improving our understanding of how global change might alter this important process of senescence. Since photoperiod is a strong driver of senescence in DB forests (32), and climatically most site-years are close to their means, the historical average is often a good predicter of the timing of senescence. However, with shifts in the timing of senescence in many populations (11), the historical average predictions are limited in their ability to predict senescence in the future under global change. Our results here emphasize the importance of understanding how global change is altering chlorophyll biosynthesis and breakdown to understand and predict changes to senescence.

We found evidence to support the transferability of model calibrations, which suggests that the model is not overfitted to the specific sites and reflects physiological mechanisms that initiate senescence. While the PhenoCam Network strives to maintain similar camera setups across their sites (33, 34), cameras and viewing angles differ across sites, which sets a high bar for assessing the transferability of model parameters, especially since we are interested in predicting greenness, not just transition dates. The significant effect of differences in peak greenness on transferability is consistent with this. Since the PhenoCam Network does not provide viewing angles in site metadata, we did not investigate this relationship here.

The two tested variables that could explain the most variation in the transferability of the calibrations, latitude and MAT, are known to affect sensitivities to drivers as plants take on different risk strategies determined by the harshness of their local winters (10). In areas with harsher winters, such as at higher latitudes or lower MATs, photoperiod is the dominant driver and in areas with milder winters, temperature is the dominant driver (10). The latitude effect on the transferability of model calibrations could also be because the shading in a PhenoCam view changes throughout the year and these changes are dependent on latitude.

### 3.2. H2: Synthesis and Breakdown Parameters Can be Estimated From Only Pre-senescence Data

Our results supported our H2 hypothesis that, if a trigger is not necessary to cause senescence in deciduous broadleaf forests, pre-senescence only data can predict the sharp decrease of greenness and the amount of curvature of inflection. That we could calibrate the model without including senescence data, nor a trigger, and still predict SOS curvature and SOS greenness values better than historical averages provides particularly strong support to our hypotheses that the SOS is initiated by unfavorable environmental conditions decreasing synthesis, and that an accumulated memory about environmental changes leading to an internal trigger is not necessary. Additionally, we found that “summer-only” calibrations can be used to predict if senescence starts sharply or more gradually, which has implications for predicting if a forest is going to rapidly or less rapidly change. This is important because the rate of senescence is likely slowing with climate change (33).

Furthermore, since we showed that the model predicted inflection in most site-years without including senescence data (Fig. S2), the ability of a pre-SOS calibration to predict greenness after SOS is not due to the influence of pre-SOS greenness. For example, a linear extrapolation of growing season greenness trends, and a standard persistence forecast (tomorrow is like today), would predict either no inflection or an inflection in the wrong direction, respectively. For the sites where pre-senescence data is insufficient to predict post-senescence greenness better than historical averages, we suspect that other environmental drivers are affecting synthesis (*e.g.,* water availability, decreased photosynthesis).

### 3.3. H3: Drivers After a Date are Better Predictor of SOS Than Those Before

Through separate random forest modeling and in support of H3, we found environmental conditions (mean temperature x mean day length) the week after a date were more important to predicting if senescence started than the environmental conditions before (Fig. 5). These results indicate that greenness can more rapidly decrease because of environmental conditions on those days, with the conditions preceding SOS less important. Therefore, predicting if post-SOS greenness values are going to rapidly decrease in the future depends on knowing (or forecasting) future environmental conditions, a feature present in our trigger-less model but absent from current CDD trigger models. Our additional results using the random forest model are consistent with the assumptions in our presented chlorophyll cycling model that assume a trigger is not necessary for senescence and that it can start as a passive process.

### 3.4. Next Steps to Improve Late Autumn Green-Down

As autumn progresses past early senescence, a range of other feedbacks and processes affect the later part of GD; therefore, it was not surprising that later parts of the GD curve could not be predicted using only pre-SOS data (Fig. 3c) and had lower transferability (Fig. S1). In these two types of validation, the rate of late-season GD was consistently too slow (*e.g.,* Fig. 2e). Since PhenoCam greenness is sensitive to leaf presence (35), the logical next step is to add in abscission by modeling the formation of the abscission layer, which due to a hormonal link with senescence constitutes a positive feedback (36). Adding a feedback between photosynthesis and chlorophyll concentrations would also cause a steeper GD because decreased photosynthesis directly decreases chlorophyll biosynthesis (19, 37–39). Implementing these processes would also prevent late autumn regreening (*e.g.,* Lac Clair, 2017 in Fig. 2c, d).

We also saw evidence of the potential to predict regreening events, but we admit it can be difficult to unambiguously distinguish regreening events from PhenoCam observational noise (Fig. 2e). To our knowledge, predicting regreening events is not possible in other senescence models. Regreening events might become more frequent as increased droughts and heat stress (40) increase summer senescence, and warmer autumns promote regreening. Thus, future work could investigate this capability to predict regreening.

### 3.5. Implications of trigger-less senescence

We have shown that senescence in DB forests may often be, at least initially, a passive process whereby synthesis is no longer able to keep pace with breakdown. This lack of a threshold-based trigger causing one-way physiological shifts in the leaves is different from what has been previously assumed (12–14, 16, 17). Studies in model herbs have shown that plant hormones are needed to trigger senescence; however, recent work has found that while the common plant hormone abscisic acid triggers senescence in model herbs, it is not essential for senescence in deciduous trees (41). We acknowledge the limitations in our study that should be investigated in future work: canopy greenness does not perfectly linearly correlate with chlorophyll concentrations; the environmental drivers of senescence are simplified here and additional drivers could be included; and our model calibrations are site-specific and are not intended to directly predict changes to senescence in the far future. Regardless of these limitations, our results and conclusions have broad impact in informing our understanding of the mechanism of senescence in DB forests.

## 4. Conclusions

Here we demonstrate that a novel mechanistic model based on chlorophyll breakdown and synthesis can predict daily phenological conditions without the need for a trigger to start senescence or a memory mechanism to keep track of cumulative environmental conditions. We show lower temperatures and photoperiods decreasing synthesis can predict the start of senescence inflection, and that in many cases such models can be calibrated successfully just based on summer data. Furthermore, we show a notable transferability of the model calibrations to other sites.

Our results are consistent with previous findings of temperature and photoperiod driving senescence and are not inconsistent with known internal physiological mechanisms. Our findings suggest that it is possible that internal transitions from anabolic to catabolic genes, which often characterizes senescence (25), could result from the decreased synthesis and other feedbacks instead of vice versa. Our model improves our understanding of the mechanisms behind canopy senescence and, thus, our ability to predict and track climate change induced changes to ecosystem functioning.

## 5. Methods

### 5.1 PhenoCam Site Selection and Data

Seventy sites were selected from the global PhenoCam Network (33, 34) as having deciduous broadleaf (DB) as a dominant forest type, representing top-of-canopy cameras, and having at least two years of low-noise data (Supplementary Table S1). Of these, twenty-four were selected for model calibration that had long PhenoCam records or represented south-temperate latitudes that did not have long records (*e.g.,* Russell Sage, Louisiana, USA had only four complete, low-noise years).

Daily PhenoCam green chromatic coordinate (*G_CC_*) 90% quantile values (33, 34) were downloaded from the PhenoCam website archive and rescaled between 0 and 1, representing winter baseline and summer peak, respectively, using the scaling factors provided by the *ElmoreFit* function in the R package ‘phenopix’ (42, 43). We filtered out years with large periods of missing data, ending up with 423 site-years across calibration and validation.

Daily photoperiods were determined as the difference between sunrise and sunset times calculated using the R package ‘suncalc’ (44). Data on temperature at 2m height was provided from the European Center for Medium-Range Weather Forecasts (ECMWF)’s hourly re-analysis product, ERA5 (45). Daily average temperature was calculated from the ERA5 data.

### 5.2. Chlorophyll Cycling Model

Daily relative canopy greenness at time *t* (*G_t_*), was modeled using a Bayesian state-space framework, which was selected because it models the latent state of a dynamic process and partitions process from observational error. In a Gaussian state space model, the latent state (*G_t_*) is dependent on the previous latent state (*G_t-1_*), a process model used to predict from time *t*-1 to time *t*, and a Gaussian process error to account for model imperfection and stochastic processes.

Our model assumes that chlorophyll breaks down at a constant exponential (23) rate, that synthesis is linearly dependent on the product of air temperature (*T_t_*) and photoperiod (*D_t_*), and that phenological greenness (*G*_t_) is positive but cannot be larger than it was at *G_t=0_* for each year. The effect of the product of air temperature and photoperiod on senescence is often included in common phenology models (12, 13). Thus, daily greenness (*G_t_*) was calculated as:

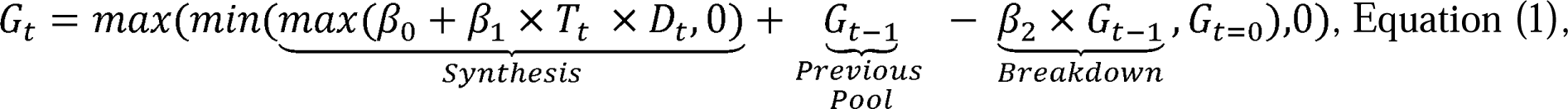

where the *β*’s represent fitted parameters. To prevent synthesis at temperatures below 0 °C and because photoperiod and temperature both positively correlate with phenological condition (11), we assigned *β*_0_, *β*_1_, and *β*_2_ uniform prior distributions *U*(*0,1*).

### 5.3. Historical Null Model

Since senescence does not usually vary greatly between years, we fit a null model that predicted that *G_t_* equals the average value on that day of year of included data plus a Gaussian process error, which was given the same prior as out chlorophyll cycling model:

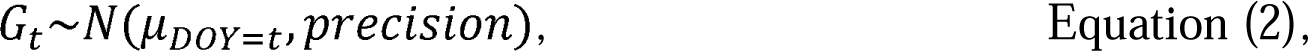

Where μ*_DOY=t_* indicates the average value of observations for that day of year.

### 5.4. Model Calibration and Validation

After randomly withholding one year per site for validation, we fit the above models separately for the twenty-four calibration PhenoCam sites using Markov Chain Monte Carlo (MCMC) in JAGS (46). JAGS was called from R version 3.5.1 (47) using the packages ‘rjags’ (48) and ‘runjags’ (49). Additional Bayesian and JAGS specific methods are given in the Supplementary Materials.

To understand how much autumn data is necessary to include for estimated parameter distributions to predict a start of senescence (SOS) inflection point, we varied the included data by setting PhenoCam observations from the period 1 August to 31 December of each year to NA (while keeping the corresponding driver data) and subsequently added in one more day of data each year to estimate new parameter distributions.

### 5.5. Model Assessment and Statistical Analyses

To determine the ability of the model to predict inflections, we calculated the differences between subsequent daily *G_CC_* values in the PhenoCam data. We then averaged these across *G_CC_* values before and after each respective date, including various amounts of observations ranging from 5 – 30 days. Next, we took the difference between the averages and concluded that the maximum inflection occurred when these averaged second differences were at a minimum. We settled on a 15-day average by visually assessing when the minimum second difference was consistently close to the day of inflection and then used this to calculate the averaged second differences of the model predictions. We also used the PhenoCam second differences to determine the minimum magnitude of second differences that constitutes an inflection point (−0.02). We then calculated the correlation between the modeled and PhenoCam peak second difference across the 24 calibration sites for various amounts of included data. We fit linear models in R using the *lm* function and investigated if the models have significantly positive slopes using the two-sided alternative hypothesis test.

To assess H1 and model performance around the SOS transition we ran a simple Bayesian model that estimated the date of the SOS changepoint between two fitted slopes (slope 1 indicating the summer greenness decline and slope 2 indicating the senescence period) after the post-senescence values were removed (*i.e.,* values less than 0.1 were removed). These models were fit as described in the above Model Calibration section. We used this methodology to estimate transition days because the second difference method was not always able to identify the precise timing of the inflection.

We scored our model predictions based on the Continuous Ranked Probability Score (CRPS), which accounts for the uncertainty in the predicted values, using *crps_sample* in the ‘scoringRules’ R package (50). We subtracted our models’ CRPS values from the historical model CRPS values across all days for all site-years to compare the models.

To answer H2, that senescence can be predicted using only pre-senescence calibration data and concurrent environmental conditions, we calculated the cumulative ratio of site-years that converged and that produced inflection points versus the number of days per year included in the calibration relative to the site’s average transition date.

To further investigate H1, for each of the 24 calibration sites, we used the model calibrated to the full autumn dataset to predict green-down at the sixty-nine other sites and then calculated the differences in CRPS compared to the validation site’s climatology for the withheld years. To investigate if there were periods of the autumn that had higher predictability than others, for each calibration-and-validation-site combination, we calculated the average CRPS difference across the full autumn, within three days of the transition, six days before the transition, and six days after the transition. Additionally, to investigate what factors affected transferability, we fit univariate regressions between the CRPS differences and the calibration and validation sites’ differences in mean annual temperature, mean total precipitation, latitude, minimum PhenoCam *G_CC_*, maximum PhenoCam *G_CC_*, average transition date, and elevation. Starting from the univariate model that had the highest coefficient of determination we created a multiple regression model via forward selection.

### 5.6. Random Forest to Assess Importance of Environmental Driver Timing on SOS

Our general hypothesis that a trigger is not necessary to start senescence also means that the inflection in declining greenness occurs because of a shift in environmental conditions and, thus, chlorophyll and greenness synthesis. If this were true then the environmental conditions right after an inflection date would have greater importance in determining if the SOS inflection occurred than conditions prior to the inflection. Alternatively, if the SOS inflection is caused by a trigger, then the conditions prior to inflection would be more important. To test this (H3), we fit a random forest model, using the R package ‘randomForest’ and the *randomForest* function’s default settings (51), that predicted if a date was the SOS inflection using the weekly mean temperature before the date, temperature after the date, day length before the date, day length after the date, temperature x day length before the date, temperature x day length after the date, site latitude, and site mean annual temperature as covariates. We randomly split the entire dataset (all 70 sites) into 75% training and 25% test. We then assessed the importance of each covariate in determining if SOS occurred or not using Mean Decrease Gini in the ‘randomForest’ R package (51).

## Author Contributions

KW and MD designed the study and KW executed the modeling and analysis. KW wrote the manuscript with inputs and suggestions from MD throughout the writing process.

## Classification

Biological sciences: ecology

## Data Availability

All data is already publicly available through PhenoCam (https://phenocam.sr.unh.edu/webcam/) or ERA5 (https://www.ecmwf.int/en/forecasts/datasets/reanalysis-datasets/era5).

## Code Availability

Code is available on KIW’s github repository: https://github.com/k-wheeler/chlorophyllCycling with information about which files are important and their file descriptions in the README.md file.

## Acknowledgements

This work was made possible by the U.S. National Science Foundation grant 1638577. KIW also acknowledges support under the NSF Graduate Research Fellowship grant 1247312 and a NOAA Climate and Global Change Postdoctoral Fellowship Program, administered by UCAR’s Cooperative Programs for the Advancement of Earth System Science (CPAESS) under award #NA21OAR4310383. Any opinion, findings, and conclusions or recommendations expressed in this material are those of the authors and do not necessarily reflect the views of the National Science Foundation. Special thanks to the Dietze lab members for feedback on the manuscript.

We thank our many collaborators, including site PIs and technicians, for their efforts in support of PhenoCam. The development of PhenoCam has been funded by the Northeastern States Research Cooperative, NSF’s Macrosystems Biology program (awards EF-1065029 and EF-1702697), and DOE’s Regional and Global Climate Modeling program (award DE-SC0016011). We acknowledge additional support from the US National Park Service Inventory and Monitoring Program and the USA National Phenology Network (grant number G10AP00129 from the United States Geological Survey), and from the USA National Phenology Network and North Central Climate Science Center (cooperative agreement number G16AC00224 from the United States Geological Survey). Additional funding, through the National Science Foundation’s LTER program, has supported research at Harvard Forest (DEB-1237491) and Bartlett Experimental Forest (DEB-1114804). We also thank the USDA Forest Service Air Resource Management program and the National Park Service Air Resources program for contributing their camera imagery to the PhenoCam archive.

## Code Availability

Code is available on KIW’s github repository https://github.com/k-wheeler/chlorophyllCycling with information about which files are important and their file descriptions in the README.md file.

