## Supplementary Materials for "A trigger may not be necessary to cause senescence in deciduous broadleaf forests"

***Supplementary Methods Related to Bayesian Modeling***

The phenological observations are related to the latent state through a Gaussian observation error model. The observation error and process error precisions were given *Gamma(shape=1.56, rate=0.016) priors*, which were selected to have a mean of 100 (*i.e.,* 0.1 standard deviation) and wide variance. Priors on *G_t=0_* for each site-year were assigned a beta distribution based on the last 10 observations for that June.

In running JAGS to calibrate the model, five chains were run to convergence (accessed using Gelman-Brooks-RubFin statistic, GBR) and until all had effective sample sizes >5000 (assessed using *rjags::effectiveSize*) after burn-in (GBR >1.05) was removed. If a model had not converged after 1,000,000 iterations, or converged to the priors, we did not include it.

In the changepoint model to determine when the SOS inflection occurred, uninformed priors of *U(-1,0)* were put on the two slope values and *U(0,100)* on the number of days after 1 August changepoint parameter.

***Figures***

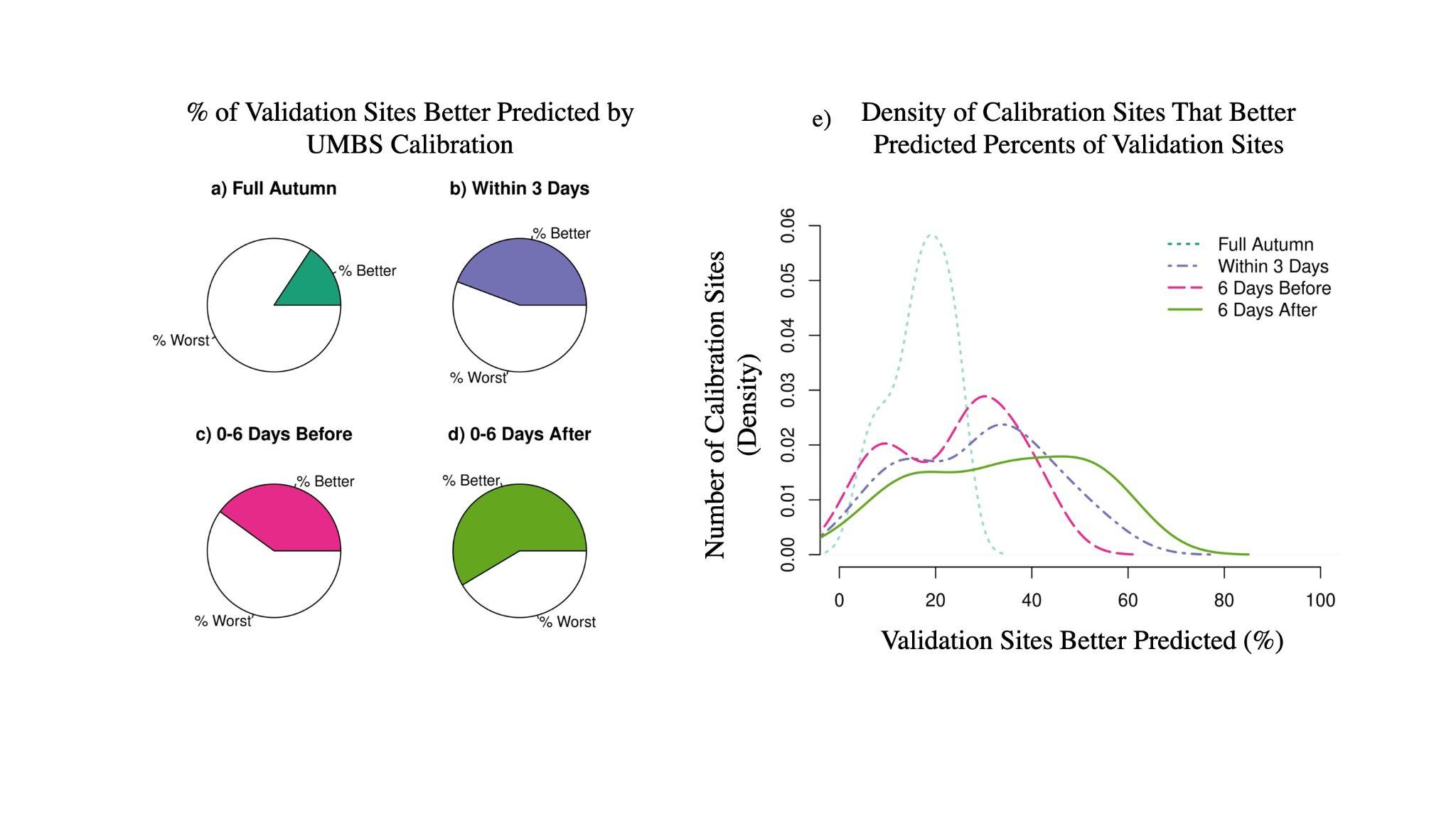

**Fig. S1.** The transferability of our model parameters for different time periods compared with the historical averages of greenness at each validation site. The percentages of validation sites that were better predicted by one example calibration of our model at University of Michigan Biological Station (UMBS) across the full autumn (a), within three days of the start of senescence (b; SOS), 0–6 days before SOS (c), and 0-6 days after SOS (d). (e) Shows how many calibration sites (*y-*axis) were able to predict greenness across the different time periods (different lines) for different percentages of validation sites (*x-*axis). The percentages in (a-d) indicate different values on the *x-*axis in (e). The model was regularly able to predict greenness around SOS at other sites better than the validation sites’ historical averages, especially during the time right after SOS.

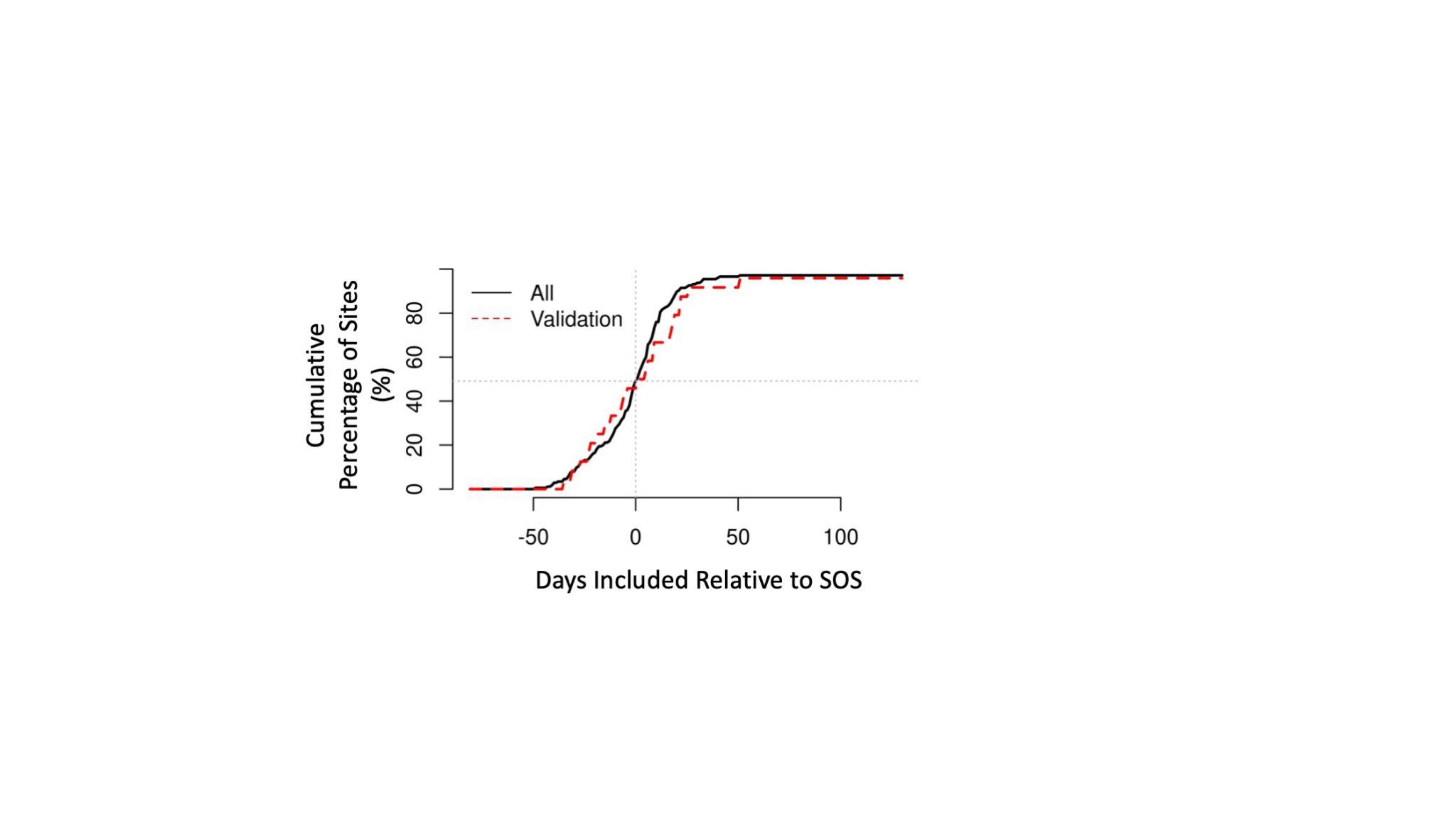

**Fig. S2.** Cumulative percentage of site-years that converged and produced an inflection based on how many days before the start of senescence (SOS) were used in the calibration data for all site-years (black solid line) and the validation years withheld from the calibration sites (red dashed line). 49% and 46% of the calibration and validation of the site-years, respectively, converged with inflection points without including senescence calibration data.

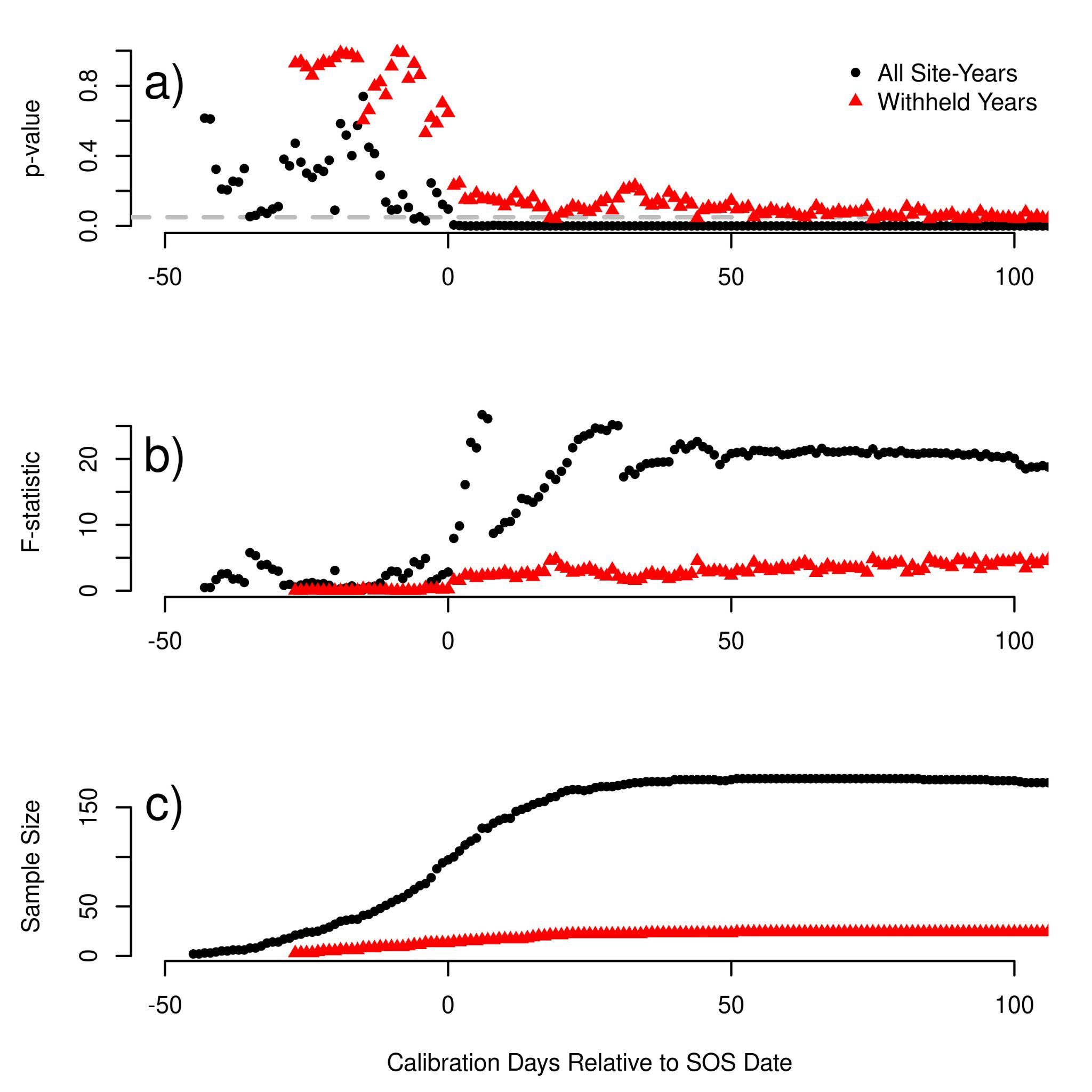

**Fig. S3.** Accompanying statistics for main text Fig. 4 for all site-years (black circles) and for the withheld years in the calibration data (red triangles). (a) depicts the change in *p*-values based on how many days of year relative to the transition date included in calibration. The gray dotted line denotes the 0.05 significant cut-off. (b) depicts the change in accompanying *F*-statistic and (c) the sample sizes. The sample sizes and power are substantially larger when including all site-years than the withheld years and larger when you include more calibration days relative to the start of senescence (SOS) date.

***Tables***

**Table S1. Selected Sites for calibration and validation**

| Site Name | Calibration Site? | # of Years | Lat | Lon | Mean SOS DOY | MAT (°C) | MAP (mm) | Elev (m) | *G_CC_* Max | *G_CC_* Min |
| --- | --- | --- | --- | --- | --- | --- | --- | --- | --- | --- |
| harvard | Yes | 12 | 42.54 | -72.17 | 281 | 6.8 | 1139 | 340 | 0.366 | 0.464 |
| proctor |  | 12 | 44.53 | -72.87 | 265 | 5 | 1081 | 403 | 0.31 | 0.357 |
| bartlettir | Yes | 11 | 44.06 | -71.29 | 255 | 5.5 | 1224 | 268 | 0.351 | 0.462 |
| umichbiological | Yes | 11 | 45.56 | -84.71 | 279 | 5.9 | 797 | 230 | 0.341 | 0.423 |
| umichbiological2 |  | 11 | 45.56 | -84.7 | 263 | 5.9 | 797 | 240 | 0.348 | 0.417 |
| bostoncommon | Yes | 10 | 42.36 | -71.06 | 292 | 9.8 | 1127 | 10 | 0.378 | 0.46 |
| harvardblo |  | 10 | 42.54 | -72.17 | 275 | 6.8 | 1139 | 340 | 0.346 | 0.435 |
| coweeta | Yes | 9 | 35.06 | -83.43 | 278 | 12.5 | 1722 | 680 | 0.357 | 0.458 |
| howland2 | Yes | 9 | 45.21 | -68.74 | 264 | 5.3 | 1058 | 79 | 0.329 | 0.423 |
| morganmonroe | Yes | 9 | 39.32 | -86.41 | 281 | 11.2 | 1087 | 275 | 0.322 | 0.382 |
| missouriozarks | Yes | 8 | 38.74 | -92.2 | 282 | 12.4 | 974 | 219 | 0.328 | 0.397 |
| queens | Yes | 8 | 44.57 | -76.32 | 254 | 6.4 | 887 | 126 | 0.325 | 0.426 |
| willowcreek | Yes | 8 | 45.81 | -90.08 | 250 | 3.9 | 820 | 521 | 0.32 | 0.436 |
| woodshole |  | 8 | 41.55 | -70.64 | 284 | 10 | 1178 | 10 | 0.37 | 0.44 |
| alligatorriver | Yes | 7 | 35.79 | -75.9 | 260 | 16.4 | 1312 | 1 | 0.328 | 0.433 |
| bostonu |  | 7 | 42.35 | -71.1 | 288 | 9.7 | 1121 | 10 | 0.306 | 0.372 |
| downerwoods | Yes | 7 | 43.08 | -87.88 | 270 | 8.2 | 812 | 213 | 0.336 | 0.431 |
| dukehw | Yes | 7 | 35.97 | -79.1 | 272 | 14.6 | 1166 | 400 | 0.314 | 0.429 |
| harvardbarn |  | 7 | 42.54 | -72.19 | 294 | 6.7 | 1151 | 350 | 0.292 | 0.389 |
| harvardbarn2 |  | 7 | 42.54 | -72.19 | 276 | 6.7 | 1151 | 350 | 0.3 | 0.42 |
| hubbardbrook | Yes | 7 | 43.94 | -71.7 | 259 | 5.6 | 1060 | 253 | 0.342 | 0.403 |
| readingma | Yes | 7 | 42.53 | -71.13 | 264 | 9.3 | 1107 | 100 | 0.31 | 0.38 |
| shalehillsczo |  | 7 | 40.67 | -77.9 | 266 | 9.8 | 980 | 310 | 0.333 | 0.414 |
| uwmfieldsta |  | 7 | 43.39 | -88.02 | 279 | 7.5 | 806 | 265 | 0.312 | 0.391 |
| ashburnham |  | 6 | 42.6 | -71.93 | 269 | 7.1 | 1147 | 292 | 0.326 | 0.38 |
| lacclair | Yes | 6 | 46.95 | -71.67 | 255 | 3.4 | 1154 | 313 | 0.307 | 0.414 |
| sanford | Yes | 6 | 42.73 | -84.46 | 272 | 8.1 | 781 | 268 | 0.307 | 0.393 |
| turkeypointdbf |  | 6 | 42.64 | -80.56 | 276 | 8 | 968 | 211 | 0.311 | 0.405 |
| worcester |  | 6 | 42.27 | -71.84 | 285 | 8.3 | 1174 | 185 | 0.329 | 0.4 |
| oakridge1 | Yes | 6 | 35.93 | -84.33 | 284 | 13.8 | 1365 | 371 | 0.331 | 0.366 |
| oakridge2 |  | 6 | 35.93 | -84.33 | 288 | 13.8 | 1365 | 371 | 0.33 | 0.374 |
| bbc1 | Yes | 5 | 42.54 | -72.17 | 293 | 6.8 | 1139 | 362 | 0.33 | 0.452 |
| bbc2 |  | 5 | 42.54 | -72.19 | 287 | 6.8 | 1139 | 380 | 0.322 | 0.467 |
| bbc5 |  | 5 | 41.55 | -70.64 | 279 | 10 | 1178 | 10 | 0.325 | 0.413 |
| bbc7 |  | 5 | 44.06 | -71.29 | 256 | 5.5 | 1224 | 289 | 0.333 | 0.443 |
| harvardlph |  | 5 | 42.54 | -72.19 | 285 | 6.8 | 1139 | 380 | 0.32 | 0.4 |
| laurentides | Yes | 5 | 45.99 | -74.01 | 262 | 3.7 | 1066 | 350 | 0.34 | 0.424 |
| ncssm |  | 5 | 36.02 | -78.92 | 304 | 14.8 | 1154 | 175 | 0.313 | 0.37 |
| springfieldma |  | 5 | 42.14 | -72.59 | 263 | 9.3 | 1115 | 56 | 0.315 | 0.371 |
| marcell |  | 5 | 47.51 | -93.47 | 263 | 2.9 | 687 | 422 | 0.322 | 0.388 |
| boundarywaters | Yes | 5 | 47.95 | -91.5 | 250 | 2.8 | 719 | 519 | 0.327 | 0.404 |
| asuhighlands |  | 4 | 36.21 | -81.7 | 267 | 10.2 | 1427 | 1016 | 0.326 | 0.41 |
| bullshoals | Yes | 4 | 36.56 | -93.07 | 276 | 13.9 | 1084 | 260 | 0.332 | 0.422 |
| canadaOA |  | 4 | 53.63 | -106.2 | 269 | 0.1 | 445 | 601 | 0.307 | 0.389 |
| millhaft |  | 4 | 52.8 | -2.3 | 282 | 9.1 | 717 | 137 | 0.36 | 0.426 |
| NEON.D01.BART.DP1.00033 |  | 4 | 44.06 | -71.29 | 246 | 5.5 | 1224 | 285 | 0.339 | 0.451 |
| NEON.D01.HARV.DP1.00033 |  | 4 | 42.54 | -72.17 | 285 | 6.8 | 1139 | 359 | 0.326 | 0.44 |
| NEON.D02.BLAN.DP1.00033 |  | 4 | 39.03 | -78.04 | 256 | 11.8 | 975 | 162 | 0.322 | 0.43 |
| NEON.D02.SCBI.DP1.00033 |  | 4 | 38.89 | -78.14 | 285 | 10.9 | 1024 | 364 | 0.324 | 0.431 |
| NEON.D05.UNDE.DP1.00033 |  | 4 | 46.23 | -89.54 | 248 | 3.7 | 846 | 529 | 0.323 | 0.422 |
| NEON.D07.ORNL.DP1.00033 |  | 4 | 35.96 | -84.28 | 277 | 13.6 | 1373 | 351 | 0.329 | 0.421 |
| NEON.D08.DELA.DP1.00033 | Yes | 4 | 32.54 | -87.8 | 291 | 17.5 | 1386 | 36 | 0.347 | 0.429 |
| russellsage | Yes | 4 | 32.46 | -91.97 | 311 | 18.1 | 1341 | 20 | 0.327 | 0.412 |
| tfforest |  | 4 | 43.11 | -70.95 | 278 | 8.1 | 1108 | 23 | 0.327 | 0.437 |
| unca |  | 4 | 35.6 | -82.55 | 295 | 12.7 | 1138 | 650 | 0.347 | 0.429 |
| silaslittle |  | 4 | 39.91 | -74.6 | 287 | 11.6 | 1128 | 33 | 0.387 | 0.574 |
| thompsonfarm2N |  | 4 | 43.11 | -70.95 | 279 | 8.1 | 1108 | 23 | 0.307 | 0.349 |
| arbutuslake |  | 3 | 43.98 | -74.23 | 257 | 4.8 | 1051 | 535 | 0.346 | 0.417 |
| macleish |  | 3 | 42.45 | -72.68 | 270 | 7.6 | 1185 | 251 | 0.333 | 0.436 |
| NEON.D02.SERC.DP1.00033 |  | 3 | 38.89 | -76.56 | 283 | 13.2 | 1068 | 30 | 0.32 | 0.419 |
| NEON.D05.STEI.DP1.00033 |  | 3 | 45.51 | -89.59 | 252 | 4.4 | 804 | 476 | 0.344 | 0.431 |
| NEON.D07.GRSM.DP1.00033 |  | 3 | 35.69 | -83.5 | 284 | 12.2 | 1385 | 589 | 0.346 | 0.442 |
| NEON.D08.LENO.DP1.00033 |  | 3 | 31.85 | -88.16 | 300 | 18 | 1499 | 10 | 0.34 | 0.407 |
| NEON.D11.CLBJ.DP1.00033 |  | 3 | 33.4 | -97.57 | 282 | 17.6 | 869 | 279 | 0.323 | 0.387 |
| nist |  | 3 | 39.14 | -77.21 | 276 | 12.1 | 1121 | 100 | 0.349 | 0.433 |
| pace |  | 3 | 37.92 | -78.27 | 271 | 13.3 | 1082 | 100 | 0.31 | 0.408 |
| robinson |  | 3 | 37.47 | -83.16 | 276 | 12.6 | 1182 | 483 | 0.349 | 0.442 |
| robinson2 |  | 3 | 37.47 | -83.16 | 279 | 12.6 | 1182 | 483 | 0.348 | 0.435 |
| morganmonroe2 |  | 2 | 39.32 | -86.41 | 280 | 11.2 | 1087 | 275 | 0.328 | 0.408 |
| NEON.D07.MLBS.DP1.00033 |  | 2 | 37.38 | -80.52 | 268 | 8.6 | 1168 | 1177 | 0.337 | 0.405 |

Note: SOS refers to start of senescence; DOY refers to day of year; mean annual temperature (MAT) and mean annual precipitation (MAP) are from World Clim^51^; *G_CC_* refers to the PhenoCam green chromatic coordinate; and Lat, Lon, and Elev refer to latitude, longitude, and elevation, respectively.

**Table S2. Calibrated parameters for each site ordered by mean *𝛽*_1_**

| Site Name | *𝛽*_0_ mean | *𝛽*_0_ sd | *𝛽*_1_ mean | *𝛽*_1_ sd | *𝛽*_2_ mean | *𝛽*_2_ sd |
| --- | --- | --- | --- | --- | --- | --- |
| bostoncommon | -0.00232 | 0.00139 | 7.00E-05 | 1.00E-05 | 0.03168 | 0.00454 |
| umichbiological | -0.01551 | 0.00335 | 0.00012 | 1.00E-05 | 0.03014 | 0.00382 |
| bbc1 | -0.01643 | 0.00463 | 0.00012 | 2.00E-05 | 0.03327 | 0.00648 |
| NEON.D08.DELA.DP1.00033 | -0.01621 | 0.00478 | 0.00012 | 2.00E-05 | 0.04582 | 0.01014 |
| dukehw | -0.02799 | 0.00406 | 0.00013 | 2.00E-05 | 0.03297 | 0.00427 |
| oakridge1 | -0.02603 | 0.00901 | 0.00013 | 3.00E-05 | 0.03127 | 0.00836 |
| russellsage | -0.01796 | 0.00422 | 0.00013 | 2.00E-05 | 0.04531 | 0.00863 |
| missouriozarks | -0.02664 | 0.0062 | 0.00014 | 2.00E-05 | 0.03367 | 0.00507 |
| bullshoals | -0.03058 | 0.00741 | 0.00014 | 3.00E-05 | 0.03944 | 0.00779 |
| sanford | -0.0235 | 0.00712 | 0.00014 | 2.00E-05 | 0.03051 | 0.00542 |
| harvard | -0.02246 | 0.00412 | 0.00017 | 2.00E-05 | 0.0384 | 0.00479 |
| readingma | -0.02819 | 0.00685 | 0.00017 | 3.00E-05 | 0.037 | 0.00603 |
| downerwoods | -0.02533 | 0.00432 | 0.00017 | 2.00E-05 | 0.04437 | 0.00628 |
| coweeta | -0.03874 | 0.00628 | 0.00021 | 3.00E-05 | 0.0489 | 0.00723 |
| queens | -0.03791 | 0.00496 | 0.00021 | 2.00E-05 | 0.05022 | 0.00595 |
| willowcreek | -0.04483 | 0.00728 | 0.00022 | 3.00E-05 | 0.039 | 0.00474 |
| bartlettir | -0.045 | 0.00651 | 0.00026 | 3.00E-05 | 0.04731 | 0.0051 |
| alligatorriver | -0.06179 | 0.00746 | 0.00026 | 3.00E-05 | 0.06478 | 0.00911 |
| lacclair | -0.04035 | 0.00683 | 0.00028 | 3.00E-05 | 0.0632 | 0.00772 |
| laurentides | -0.06253 | 0.02504 | 0.00028 | 7.00E-05 | 0.03327 | 0.00769 |
| boundarywaters | -0.06181 | 0.01506 | 0.00028 | 5.00E-05 | 0.03529 | 0.00621 |
| morganmonroe | -0.06144 | 0.02784 | 0.00031 | 0.00011 | 0.02806 | 0.00582 |
| howland2 | -0.08186 | 0.02787 | 4.00E-04 | 0.00011 | 0.04023 | 0.00749 |

Note: sd refers to standard deviation.

**Table S3. Percentage of withheld site-years for each calibration site better predicted than historical averages of validation sites**

| **Calibration Site** | **% Better 0 – 6 Days After** | **% Better Within 3 Days** | **% Better 0 – 6 Days Before** | **% Better Full Autumn** |
| --- | --- | --- | --- | --- |
| bostoncommon | 60 | 56 | 36 | 6 |
| umichbiological | 59 | 44 | 40 | 16 |
| harvard | 56 | 44 | 43 | 17 |
| morganmonroe | 54 | 53 | 41 | 14 |
| hubbardbrook | 51 | 46 | 39 | 17 |
| sanford | 51 | 41 | 34 | 19 |
| bbc1 | 49 | 39 | 29 | 19 |
| howland2 | 46 | 33 | 29 | 20 |
| readingma | 46 | 36 | 33 | 20 |
| laurentides | 40 | 31 | 30 | 23 |
| boundarywaters | 36 | 31 | 30 | 23 |
| lacclair | 34 | 31 | 26 | 24 |
| bartlettir | 34 | 31 | 27 | 26 |
| downerwoods | 34 | 31 | 26 | 24 |
| willowcreek | 33 | 30 | 26 | 23 |
| missouriozarks | 24 | 17 | 16 | 16 |
| oakridge1 | 24 | 19 | 13 | 14 |
| russellsage | 17 | 13 | 9 | 11 |
| coweeta | 16 | 16 | 14 | 23 |
| queens | 14 | 17 | 10 | 19 |
| dukehw | 14 | 10 | 7 | 10 |
| NEON.D08.DELA.DP1.00033 | 13 | 10 | 7 | 7 |
| bullshoals | 9 | 6 | 4 | 9 |
| alligatorriver | 1 | 3 | 3 | 4 |

Note: For different periods, what percentage of the withheld years at other sites was each calibration site better at predicting than the validation site’s historical averages.
